# Direct and indirect effects of high temperatures on fledging success in a cooperatively breeding bird

**DOI:** 10.1101/2021.01.24.427934

**Authors:** Amanda R. Bourne, Amanda R. Ridley, Claire N. Spottiswoode, Susan J. Cunningham

## Abstract

High temperatures and low rainfall consistently constrain reproduction in arid-zone bird species. Understanding the mechanisms underlying this pattern is critical for predicting how climate change will influence population persistence and to inform conservation and management. In this study, we analysed Southern Pied Babbler *Turdoides* nestling survival, daily growth rate and adult investment behaviour during the nestling period over three austral summer breeding seasons. High temperatures were associated with lower body mass, shorter tarsi, and reduced daily growth rates of nestlings. Piecewise structural equation modelling suggests that direct impacts of temperature had the strongest influence, followed by changes in provisioning rates by adults to older nestlings.

Adjustments to adult provisioning strategies did not compensate for direct negative effects of high air temperatures on nestling body mass, tarsus and wing length, or daily growth rates. Declining reproductive success during hot weather poses a potentially serious threat to population replacement and persistence as climate change progresses, even for currently common and widespread species. Significantly lower offspring survival as a result of climate warming will likely contribute to the collapse of animal communities well before temperatures regularly approach or exceed lethal tolerance limits. Detailed mechanistic data like these allow us to model the pathways by which high temperature causes nest failure. In turn, this could allow us to design targeted conservation action to effectively mitigate climate effects.

## Introduction

Anthropogenic climate change is affecting wildlife populations around the world (Saino et al. 2011; Iknayan and Beissinger 2018; Ripple et al. 2019; Rosenberg et al. 2019; van de Ven, McKechnie, et al. 2020), in part via impacts of altered temperature and rainfall patterns on reproductive success (Stevenson and Bryant 2000; Cahill et al. 2013; Cunningham et al. 2013; Paniw et al. 2019; van de Ven, Fuller, et al. 2020). Accurately predicting how climate change will influence reproduction, and hence population persistence, requires an understanding of the mechanistic links between climate and reproductive outcomes (Conradie et al. 2019; Ratnayake et al. 2019). Therefore, we need to understand the mechanisms by which temperature and rainfall affect reproductive outcomes via direct effects on offspring size, daily growth rates, and survival, as well as indirectly via effects on parental care strategies (Buchholz et al. 2019; van de Ven, McKechnie, et al. 2020).

Adverse weather conditions impair nestling development (Mainwaring et al. 2016; Imlay et al. 2018) by causing a trade-off between devoting energy to thermoregulation or to growth (Dawson et al. 2005). High temperatures constrain nestling growth (Cunningham et al. 2013; Mainwaring and Hartley 2016; Andreasson et al. 2018), result in smaller nestlings overall (Salaberria et al. 2014; Wada et al. 2015; Rodriguez and Barba 2016), alter corticosterone levels (Newberry and Swanson 2018; Crino et al. 2020) and reduce survival probabilities (Greño et al. 2008; Zuckerberg et al. 2018; Bourne, Cunningham, et al. 2020a). Higher rainfall often has a positive effect on nestling development (Wiley and Ridley 2016) and survival (Skagen and Yackel Adams 2012; Mares et al. 2017), at least in arid and semi-arid ecosystems (Cumming and Bernard 1997; Hidalgo Aranzamendi et al. 2019), although see Morganti et al. (2017) and Cox et al. (2019) for effects of rainy weather in temperate environments. Nestlings are substantially less likely to survive during droughts compared to wetter periods (Morrison and Bolger 2002; Conrey et al. 2016; Cruz-McDonnell and Wolf 2016; Bourne, Cunningham, et al. 2020b).

Effects of temperature and rainfall on nestlings may be direct, due to impacts on nestling physiology, or indirect, via for example impacts on parental care behaviours and investment strategies (Drent and Daan 1980; Salaberria et al. 2014; van de Ven, McKechnie, et al. 2020). Several recent studies suggest that negative effects of adverse weather can be moderated by adjustments in parental care strategies including brooding (Oswald et al. 2008; Mainwaring and Hartley 2016) and provisioning (Auer and Martin 2017; Sofaer et al. 2018). Other studies indicate that birds trade off foraging behaviour against thermoregulation when provisioning nests, reducing provisioning rates as temperatures rise (Cunningham et al. 2013; Wiley and Ridley 2016; van de Ven et al. 2019).

The Southern Pied Babbler *Turdoides bicolor* (hereafter pied babbler) is a medium sized (60-90 g) cooperatively breeding passerine endemic to a semi-arid ecosystem, the Kalahari Desert, characterised by hot summers and periodic droughts (Hockey et al. 2005; van Wilgen et al. 2016). Rainfall is extremely variable between years (MacKellar et al. 2014) and, over the last 20 years, high temperature extremes within the Kalahari have increased in both frequency and severity (Kruger and Sekele 2013; Bourne, Cunningham, et al. 2020a). Previous research on this species has shown that high air temperatures during early development are associated with reduced provisioning rates to nestlings (Wiley and Ridley 2016), smaller nestlings (Wiley and Ridley 2016), reduced likelihood of fledging at least one chick per breeding attempt (Bourne, Cunningham, et al. 2020a), lower recruitment of young into the adult population (Bourne, Cunningham, et al. 2020c) and compromised adult foraging efficiency and body mass maintenance (du Plessis et al. 2012). In short, pied babblers at our study site are likely to reproduce less successfully during droughts (Bourne, Cunningham, et al. 2020b) and completely fail to fledge young at mean daily maximum air temperatures of > 38°C between hatching and fledging (Bourne, Cunningham, et al. 2020a).

Pied babblers are territorial year-round residents and a habituated population at the study site enables collecting detailed life history and behavioural data from known individuals (Ridley 2016). Here, we use detailed nestling morphometric data and adult behavioural data collected from marked individuals to explore the mechanisms by which temperature, rainfall and group size influence the growth and survival of young from hatching to fledging. Specifically, we used piecewise structural equation modelling [piecewise SEM, (Shipley 2009; Larson et al. 2015; van de Ven, McKechnie, et al. 2020)] to empirically test whether temperature, rainfall, and group size influence fledging probabilities via direct effects on nestling mass and structural size (suggesting a physiological mechanism) or via the indirect effects of adjustments in adult investment behaviour (suggesting a behavioural mechanism). We further explored temperature, rainfall and group size effects on nestling daily growth rates and the foraging and provisioning behaviour of adult group members, both important for interpreting components of the piecewise SEM analyses. Detailed mechanistic data like these allow us to model the pathways by which high temperatures result in nest failure. In turn, this could allow us to design targeted conservation action to effectively mitigate climate effects.

Building on previous work demonstrating the impact of temperature on nestling body mass and adult provisioning rates in pied babblers (Wiley and Ridley 2016), we tested the hypothesis that high temperatures and low rainfall should reduce nestling size and growth via a combination of both direct effects on nestlings and indirect effects via reduced provisioning effort. Because pied babblers are cooperative breeders that engage in load-lightening behaviours (Raihani and Ridley 2008; Ridley and Raihani 2008; Wiley and Ridley 2016), we further tested the hypothesis that individuals in larger groups should allocate more time to self-maintenance activities (such as foraging, resting and preening) during hot weather while maintaining the overall biomass provisioned to young, sustaining nestling growth rates.

## Materials and Methods

### Study site and system

Data were collected for each austral summer breeding season between September 2016 and February 2019 (three breeding seasons in total) at the Kuruman River Reserve (33 km^2^, KRR; 26°58’S, 21°49’E) in the southern Kalahari. Mean summer daily maximum temperatures in the region average 34.7 ± 9.7 °C and mean annual precipitation averages 186.2 ± 87.5 mm (1995–2015, van de Ven, McKechnie & Cunningham 2019). The Kalahari region is characterised by hot summers and periodic droughts (van Wilgen et al. 2016), with extremely variable rainfall between years (MacKellar et al. 2014). Pied babblers live in groups ranging in size from 3–15 adults (Raihani and Ridley 2007a) and breed during the austral summer (September–March), when it is hottest (Ridley 2016). Pied babbler groups consist of a single breeding pair with subordinate helpers of both sexes (Nelson-Flower et al. 2011), and all adult group members (individuals > 1 year old) engage in cooperative behaviours including territory defence and parental care (Ridley and Raihani 2007; Ridley 2016). Pied babblers lay clutches of ~3 eggs (Ridley 2016), which hatch after 14 ± 1.2 days (Bourne, Cunningham, et al. 2020a), and nestlings fledge after 15.4 ± 1.7 days (Bourne, Cunningham, et al. 2020a).

### Nestling size and daily growth rates (nestling age day 5 and day 11)

Following Ridley and van den Heuvel (2012), we monitored all nests initiated (clutches laid and incubated) in the study population during each breeding season to determine hatching dates (*n* = 103 nests in total; 2016/17 *n* = 61, 2017/18 *n* = 22, 2018/19 *n* = 20). Nestlings were measured (body mass, tarsus length, and wing length) between 06h00 and 07h00 (morning) and again between 18h00 and 19h00 (evening) on the 5^th^ day after hatching (d5, representing growth during a fast growth phase; *n* = 93 nestlings from 37 broods) and the 11^th^ day after hatching (d11, representing growth during an asymptote phase; *n* = 77 nestlings from 34 broods). Body mass measurements (± 0.1 g) were taken by weighing nestlings on a top-pan scale. Tarsus length was measured (± 0.1 mm) using clock-dial Vernier calipers and wing length (± 0.1 mm) using a stopped rule. All nestling measurements were taken by the same person. Data are presented for right tarsus and right wing. Natal group size (the number of adults present during the period between hatching and fledging) and brood size (the number of nestlings in the brood on each measurement day) were recorded for each brood on each measurement day.

Nestling size was recorded as evening body mass, tarsus length and wing length, representing nestling mass and size at the end of a full day of provisioning by adults. Nestling daily growth rates were calculated as the percentage change (Δ) in body mass, tarsus length, and wing length between morning and evening measurements, standardised for differences in the time between measurements using the equation presented by du Plessis et al (2012):

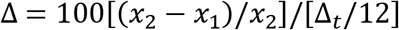

where Δ_t_ = number of hours between *t*_*1*_ (time of morning measurement) and *t*_*2*_ (time of evening measurement); *x*_*1*_ = mass, tarsus, or wing length measurement at *t*_*1*_ and *x*_*2*_ = measurement at *t*_*2*_.

### Provisioning rates to nests (nestling age day 5 and day 11)

We recorded the number of provisioning visits to the nest per unit time observed during the morning (08h00 to 09h30), at midday (12h00 to 13h00), and in the afternoon (15h00 to 16h30) on d5 (*n* = 26 broods) and d11 (*n* = 25 broods), between morning and evening nestling measurements. Data on provisioning visits were collected using a combination of video cameras (Sony HDR-XR160E) placed on a tripod 4–6 m from the nest and nest watches undertaken by one human observer with binoculars, seated 15–20 m from the nest. Provisioning data were captured using CyberTracker software (v3.448; www.cybertracker.org) on an Android smartphone (Mobicel *TRENDY*).

### Adult behaviour data (nestling age day 7-9)

To determine effects of temperature, rainfall and group size on the proportion of time adult birds allocated to parental care vs. self-maintenance, we conducted 20-minute continuous time-activity focal behaviour observations (Altmann 1974) on up to four different adult birds per day within each of 6 focal sessions. We focused on groups with 7- to 9-day-old nestlings, i.e. between the days on which we had collected nestling morphometric measurements, at 26 different broods. Focal sessions lasted two hours each, with the first starting at 07h00 and the last starting at 17h00 (i.e. 07h00 to 08h59, 09h00 to 10h59, 11h00 to 12h59, 13h00 to 14h59, 15h00 to 16h59, 17h00 to 19h00). We collected 593 focal behavioural observations (mean focal length = 19 ± 1.9 min) over 108 focal days (mean daily observation length over a focal day = 108 ± 17 min) during three austral summer breeding seasons (2016/17 *n* = 60 days, 2017/18 *n* = 32 days, 2018/19 *n* = 16 days). We observed 29 males, 29 females, and 4 individuals of unknown sex. Of these, 27 were dominant individuals and 35 were subordinates. When group size was = 3 adult individuals, the pair and the single adult subordinate were studied; when group size was = 4, the pair and both subordinate adults were studied; when group size was > 4, the pair and two subordinate adults of opposite sex (where possible) were studied. We observed each focal individual once within each of the six daily focal sessions, and randomised the order in which each individual was observed within each focal session (*sensu* du Plessis et al (2012)). In this way, we collected approximately six focal behaviour observations per bird per day, spread evenly across the day to minimise time of day effects on summarised measurements. All birds for which we had fewer than four focal observations per day were removed from the analyses (*n* = 6 of 68 individuals). From these data, we could estimate individual investment in young at the nest (including number of provisioning visits to the nest, biomass caught vs. provisioned, and time spent attending the nest for each individual), but not the total number of provisions made to the nest by all group members on that day (because we had to concentrate on one focal bird at a time). Behaviour data were captured on the Mobicel smartphone using Prim8 software (McDonald and Johnson 2014), in which the duration of each observed behaviour can be recorded to the nearest second.

For analyses of time budgets, we summed time observed foraging (foraging effort, including searching for and handling prey), attending the nest (all visits to the nest, including provisioning, shading and brooding), resting (preening, standing, and perching) and engaging in other activities (e.g. walking, flying, on sentinel duty, interacting with neighbouring groups) and calculated the proportion of time allocated to each set of activities across all six focals, at the scale of a ‘focal day’. During each focal behaviour observation, we collected detailed information for each successful foraging event, including the size class of each item caught and whether or not the item was provisioned to the nest. We converted prey captures to biomass (wet g) using the calculations from Raihani & Ridley (2007b). We recorded foraging success as total biomass caught per bird per focal day, and provisioning rates to the nest as total biomass provisioned per bird per focal day. Time allocations to different behaviours were averaged over all focals per individual as focal observations were spread evenly across the day to minimise time-of-day effects on behavioural investment patterns between individuals (*F*_*5,489*_ = 1.283, *P* = 0.269; Fig S1) and ensure that data analysed at the scale of the focal day were comparable between birds and days.

### Temperature and rainfall

Daily maximum temperature (T_max_, in °C) and rainfall (mm) data were collected from an on-site weather station at the KRR (Vantage Pro2, Davis Instruments, Hayward, CA, USA). Weather variables included in statistical models were T_max_ on the measurement day and rainfall summed for the 60 days prior to initiation of the breeding attempt (Rain_60_), to allow for known delays between rainfall and invertebrate emergence in the Kalahari (Cumming & Bernard, 1997; Ridley & Child, 2009).

#### Statistical analyses

Statistical analyses were conducted in the R statistical environment, v 3.6.0, using R Studio (R Core Team 2017).

### Nestling mass, size and survival (nestling age day 5 and day 11)

In other bird species, larger nestlings with longer tarsi and more developed wings are more likely to survive to fledging (Kruuk et al. 2015; Mumme et al. 2015; Martin et al. 2018), and prior research on pied babblers has shown that nestling mass is influenced by environmental factors such as temperature and rainfall (Wiley and Ridley 2016). We therefore undertook a series of piecewise SEM analyses (Shipley 2009; Larson et al. 2015; van de Ven, McKechnie, et al. 2020) to specify and simultaneously quantify all hypothesised relationships regarding whether and to what extent the impacts of weather conditions on survival to fledging, via d5 and d11 evening body mass, tarsus length, and wing length, are direct (i.e. could be inferred to result from physiological limitations of nestlings) or indirect (i.e. mediated via changes in adult behaviour and therefore provisioning rates to nestlings). We computed piecewise SEMs using the R package *piecewiseSEM* (Lefcheck 2016), which can accommodate multiple error structures. This capacity is important because the response terms of our component models have different distributions (see below). Path coefficients are partial regression coefficients and can be interpreted similarly to simple and multiple regression outputs. Unstandardised effect sizes are reported and statistical significance taken as *P* < 0.05 (Lefcheck 2016).

We used piecewise SEM analyses to test the following statistical hypotheses:

Survival to fledging is negatively affected by low nestling body mass, short tarsi, and short wings (logit);
Nestling body mass, tarsus length, and wing length are negatively affected by a) high T_max_ on the measurement day, b) low Rain_60_, c) smaller group size, d) larger brood size, and e) fewer provisioning events (Gaussian);
Number of provisioning events is negatively affected by a) high T_max_, b) low Rain_60_, c) smaller group size, and d) smaller brood sizes (Poisson).

### Nestling daily growth rates (nestling age day 5 and day 11)

Because daily growth rates influence the body mass, tarsus length, and wing length attained by d5 and d11 nestlings at the time of the evening measurement, we considered the effect of Rain_60_, T_max_, brood size, and natal group size on the percentage change (Δ) in body mass, tarsus length, and wing length between morning and evening measurements on d5 and d11, including brood identity as a random term. The inclusiuon of group identity as a random terms in additon to brood identity resulted in unstable models and of the two random terms, brood identity explained the greatest proportion of variation while avoiding destablising the models (Grueber et al. 2011; Harrison et al. 2018a). We used maximum likelihood linear mixed-effects models (LMMs) with Gaussian error structure in the R package *lme4* (Bates et al. 2015).

Model selection using the Akaike’s information criterion corrected for small sample size (AICc) was used to determine the model/s that best explained patterns of variation in the data. Lower AICc values were taken to represent more parsimonious models, following Harrison et al (2018b). Where there were several models within 2 AICc of the top model, top model sets were averaged using the package *MuMIn* (Barton 2015). Model estimates with confidence intervals that did not intersect zero were considered to explain significant patterns in the data (Grueber et al. 2011). Model fits were evaluated using histograms of residuals and Normal Q-Q plots.

### Adult behaviour (nestling age day 7-9)

To determine which variables best predicted the proportion of time adults spent foraging, resting, and attending the nest, we fitted binomial GLMMs with Penalised Quasi-Likelihood (glmmPQL) in the R package *MASS* (Venables and Ripley 2002). The glmmPQL approach, which precludes model selection (Bolker et al. 2009), was used to address overdispersion in the data not adequately resolved by the inclusion of an observation level random term while still allowing inclusion of the random term for brood identity. Proportion of time spent foraging was modelled as a combined vector of total time spent on the selected activity (seconds) versus total time observed (seconds). The models included predictor variables T_max_, Rain_60_, group size, and brood size, as well as sex and rank of the focal bird. Nestling age was not included in the models because all observations were collected when nestlings were 7-9 days old.

To determine which variables best predicted biomass caught and biomass provisioned, we fitted GLMMs with a Poisson error distribution (log link) in the package *lme4* (Bates et al. 2015). Model selection was undertaken as described for nestling daily growth rates above. Response variables were rounded to the nearest digit (biomass in g). We considered the influence of T_max_, Rain_60_, group size, brood size, sex, and rank, and included quadratic terms for T_max_ and group size when there was no significant linear effect and visualisation of the data suggested a non-linear relationship. Model estimates with confidence intervals that did not intersect zero were considered to explain significant patterns in the data (Grueber et al. 2011). Model fits were evaluated using the dispersion parameter. Nestling age was not included in the models because all observations were collected when nestlings were 7-9 days old. Biomass was modelled as the total amount per observation day because fitting these data resulted in models that were more stable than those generated by fitting rates (e.g. biomass per hour).

## Results

Maximum temperatures on data collection days ranged from 27.2 °C to 41.6 °C (mean = 33.7 ± 3.5 °C), and Rain_60_ ranged from 0.8 to 174.2 mm (mean = 60.2 ± 55.5 mm). Group sizes averaged 4 ± 1 adults (range: 2 to 6) and brood sizes averaged 3 ± 1 nestlings (range: 1 to 5). Of 99 monitored nests, 65 clutches hatched, 52 broods survived to 5-days-old (first nestling measurement day), 41 broods survived to 7- to 9-days-old (adult behaviour observation day), and 38 broods survived to 11-days-old (second nestling measurement day). We did not collect all data types at every breeding attempt and sample sizes achieved are stated for each analysis below.

### Nestling mass, size, and survival (nestling age d5 and d11)

Piecewise SEM analyses for d5 and d11 nestling mass explained 11% and 22% of the observed variation in fledging probability respectively (d5: *Fisher’s C* for tests of directed separation = 11.95, *df* = 10, *P* = 0.288, Fig. 1; d11: *Fisher’s C* = 15.78, *df* = 10, *P* = 0.106, Fig. 2). Piecewise SEM analysis showed that, for d5 nestlings, direct effects of temperature on the measurement day had a significant influence on nestling mass and therefore the probability of surviving until fledging, whereas indirect effects mediated via provisioning rate did not (Fig 1). In contrast, both of these pathways were statistically significant for d11 nestings (Fig 2).

**Figure 1:**
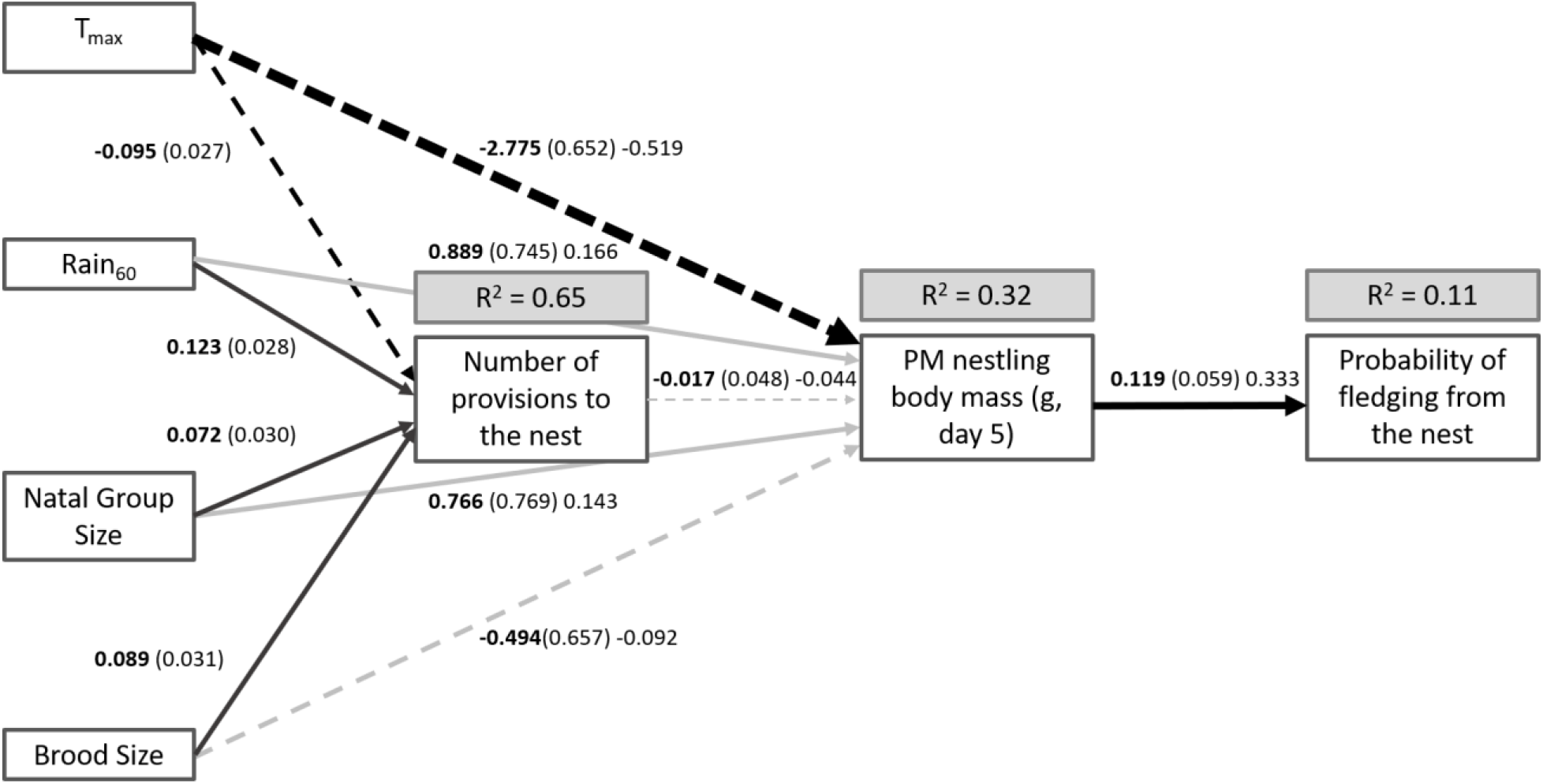
Piecewise SEM exploring the effects of environmental factors (temperature and rainfall), group size, and brood size on individual probabilities of fledging via number of provisioning visits and nestling mass (evening) 5 days after hatching. Boxes represent measured variables. Arrows represent unidirectional relationships among variables. Solid arrows denote positive relationships, dashed arrows negative relationships. Path coefficients are shown in bold, followed by standard errors in parentheses. Standardised coefficients are shown after standard errors for all pathways except those in the Poisson-distributed model with number of provisions to the nest as the response variable, for which standardised estimates could not be calculated. Non-significant paths are grey. Path thickness has been scaled relative to the absolute magnitude of the standardised estimates, such that stronger effects have thicker arrows. R^*2*^ for component models are given in the grey boxes above response variables.

**Figure 2:**
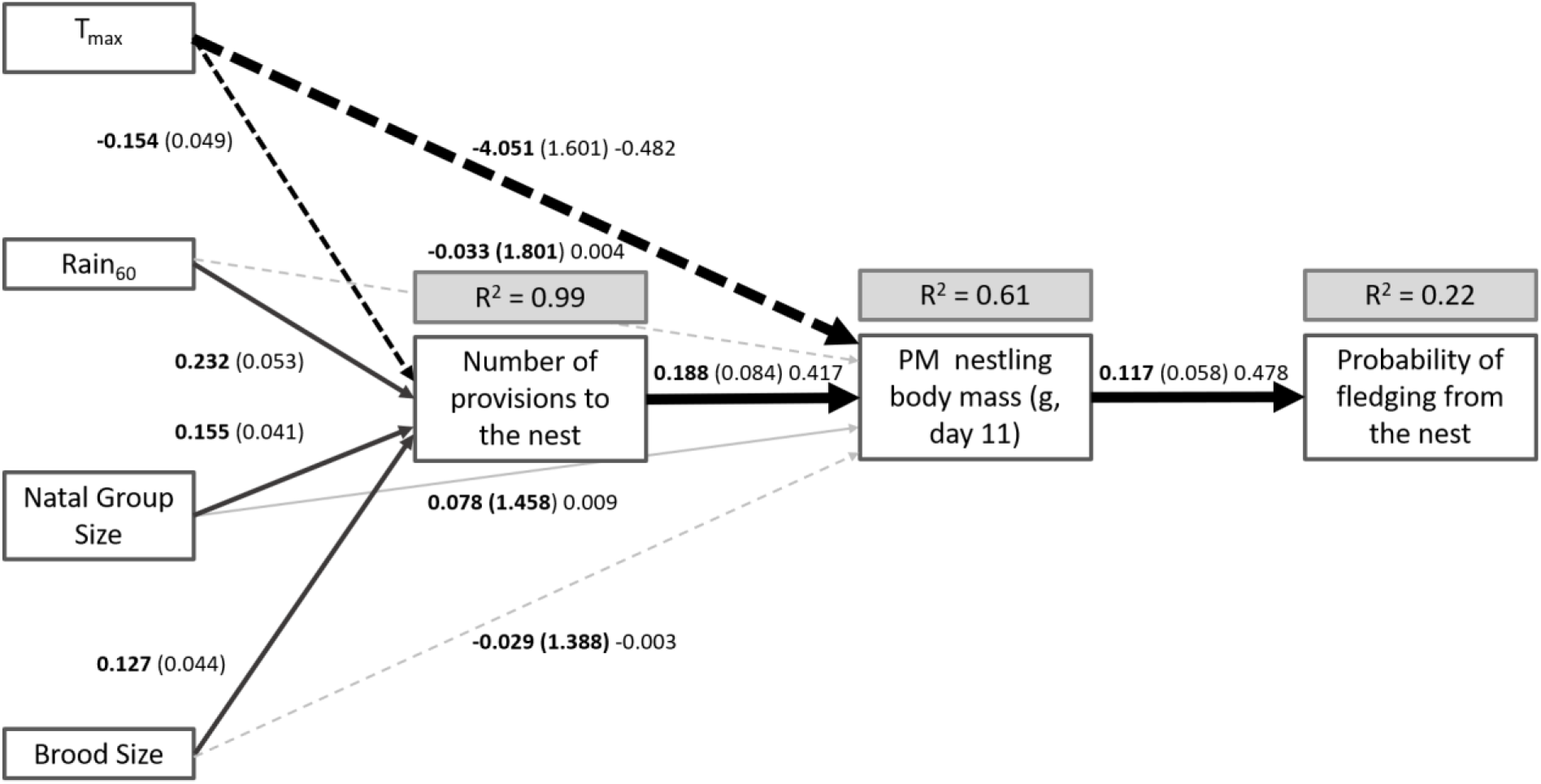
Piecewise SEM exploring the effects of environmental factors (temperature and rainfall), group size, and brood size on individual probabilities of fledging via number of provisioning visits and nestling mass (evening) 11 days after hatching. Boxes represent measured variables. Arrows represent unidirectional relationships among variables. Solid arrows denote positive relationships, dashed arrows negative relationships Path coefficients are shown in bold, followed by standard errors in parentheses. Standardised coefficients are shown after standard errors for all pathways except those in the Poisson-distributed model with number of provisions to the nest as the response variable, for which standardised estimates could not be calculated. Non-significant paths are grey. Path thickness has been scaled relative to the absolute magnitude of the standardised estimates, such that stronger effects have thicker arrows. R^*2*^ for component models are given in the grey boxes above response variables.

Specifically, high temperatures were directly associated with smaller nestling body mass on both d5 and d11 (d5: Est = −2.775, *P* < 0.001; d11: Est = −0.482, *P* = 0.018), which in turn predicted the probability of surviving to fledging in both cases (d5: Est = 0.119, *P* = 0.042; d11: Est = 0.478, *P* = 0.043). Additionally, for both d5 and d11 nestlings, provisioning efforts by adults were negatively associated with high temperatures (d5: Est = −0.095, *P* < 0.001; d11: Est = −0.154, *P* = 0.002). Number of provisioning visits to the nest averaged 31 on cool days (T_max_ < 35.5°C) and 22 on hot days (T_max_ ≥ 35.5°C) for d5 nestlings, and 42 and 26 respectively for d11 nestlings. For both d5 and d11 nestlings, provisioning efforts by adults were positively associated with Rain_60_ (d5: Est = 0.123, *P* < 0.001; d11: Est = 0.232, *P* < 0.001), group size (d5: Est = 0.089, *P* = 0.018; d11: Est = 0.155, *P* < 0.001), and brood size (d5: Est = 0.089, *P* = 0.004; d11: Est = 0.127, *P* = 0.004). Number of provisioning visits to the nest predicted larger nestling mass (and therefore higher probability of fledging) on d11 only (d5: Est = −0.017, *P* = 0.726, d11: Est = 0.417, *P* = 0.033).

Piecewise SEM for tarsus length and wing length explained < 4% of the variation in fledging probability in d5 nestlings and 33–36% of the variation in fledging probability in d11 nestlings (*Fisher’s C* < 15.08, *df* = 10, *P* > 0.129 in all cases, Appendix Fig. S2–S5). Relationships between T_max_, group size, Rain_60_, brood size and provisioning rates were the same as for nestling mass models above as these used identical datasets. Neither tarsus nor wing length on d5 influenced fledging probabilities, but nestlings with longer tarsi (Est = 0.510, *P* = 0.016) and longer wings (Est = 0.213, *P* = 0.021) on d11 were more likely to fledge. We found no evidence that tarsus or wing length were themselves influenced by T_max_, Rain_60_, group size, brood size, or number of provisioning visits for either nestling age class.

### Nestling daily growth rates (nestling age d5 and d11)

T_max_ was the most parsimonious predictor of daily mass gain both in d5 and d11 nestlings, consistent with the results of the piecewise SEM (i.e. that T_max_ was an important predictor of evening body mass). Nestlings gained less mass, and sometimes even lost mass, between morning and evening measurements on hotter days. For both d5 and d11 nestlings, we found single best-fit models for daily mass change containing only T_max_ (d5: model weight = 0.645; *n* = 93; Est = − 4.043 ± 1.986, 95% CI: −7.922, −0.151, *t* = −2.031; Fig. 3A; d11: model weight = 1.00; *n* = 77; Est = −7.028 ± 1.122, 95% CI: −8.393, 4.034, *t* = −6.262, Fig. 3B). Percentage change in tarsus length and wing length over the same 12 h period was not influenced by any of the included predictor variables in d5 nestlings, but nestling tarsi grew significantly less between morning and evening measurements on hotter days in d11 nestlings (top model weight = 0.772; Est = −0.804 ± 0.287, 95% CI: −1.381, −0.285, *t* = −2.797; Fig. 3C). Although the effect was not statistically significant, d11 nestling wings also tended to grow less on hotter days (Est = −0.651 ± 0.396, 95% CI: −1.427, 0.125, *t* = −1.643; Fig. 4D). We found no evidence for effects of Rain_60_, brood size, or natal group size in any of the nestling daily growth rate analyses (see Table S1–S3 for full model selection outputs).

**Figure 3:**
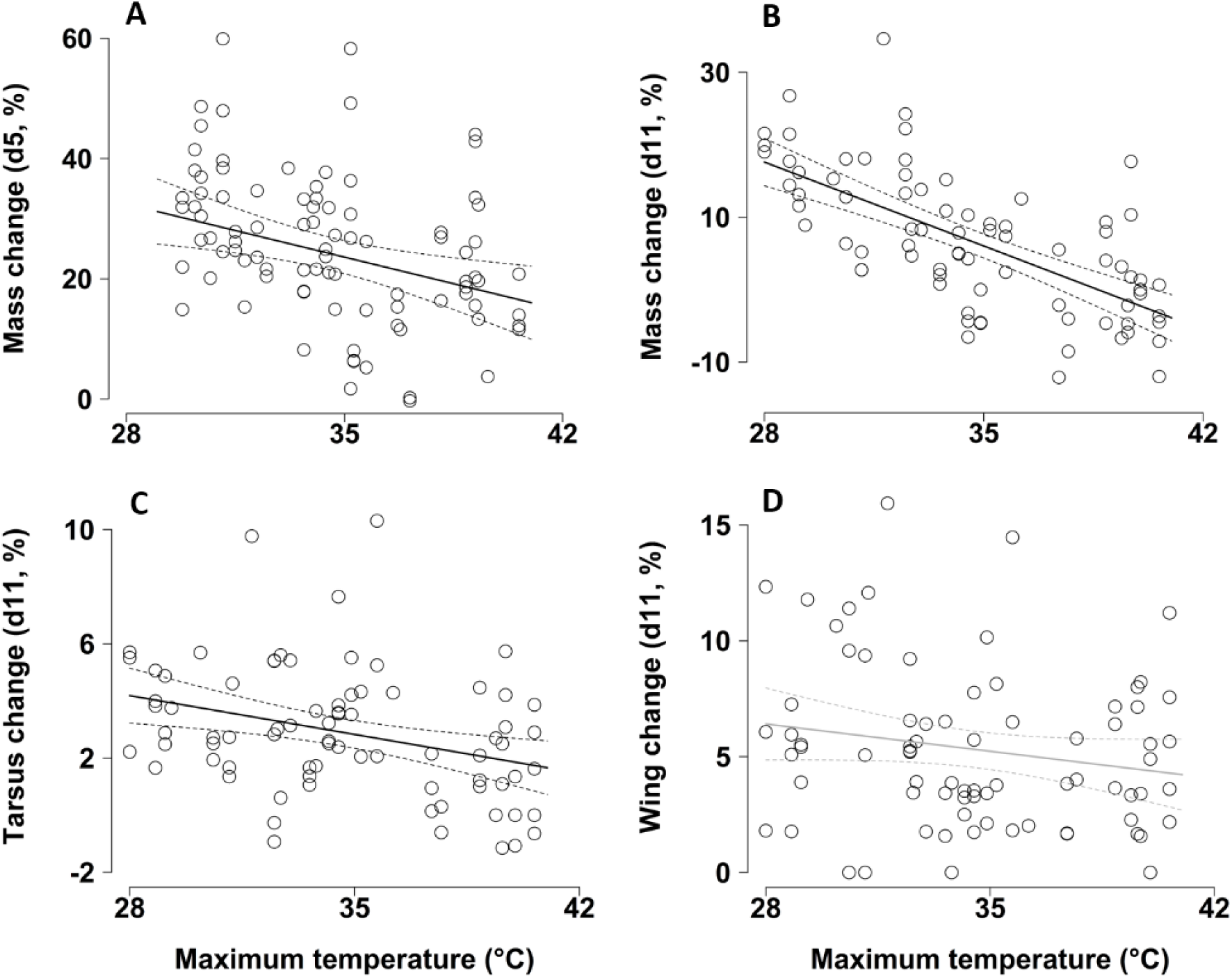
The effect of maximum daily temperature (T_*max*_, °C) on nestling daily growth rates (% change over a 12 h period). Data points show the % daily mass change of (A) 5-day-old southern pied babbler Turdoides bicolor nestlings and (B) 11-day-old nestlings as well as the (C) % daily tarsus length change and (D) wing length change of 11-day-old nestlings. Solid lines represent predictions from the models and dashed lines the 95% CIs. The regression line in (D) is greyed out as the trend shown was not statistically significant.

**Figure 4:**
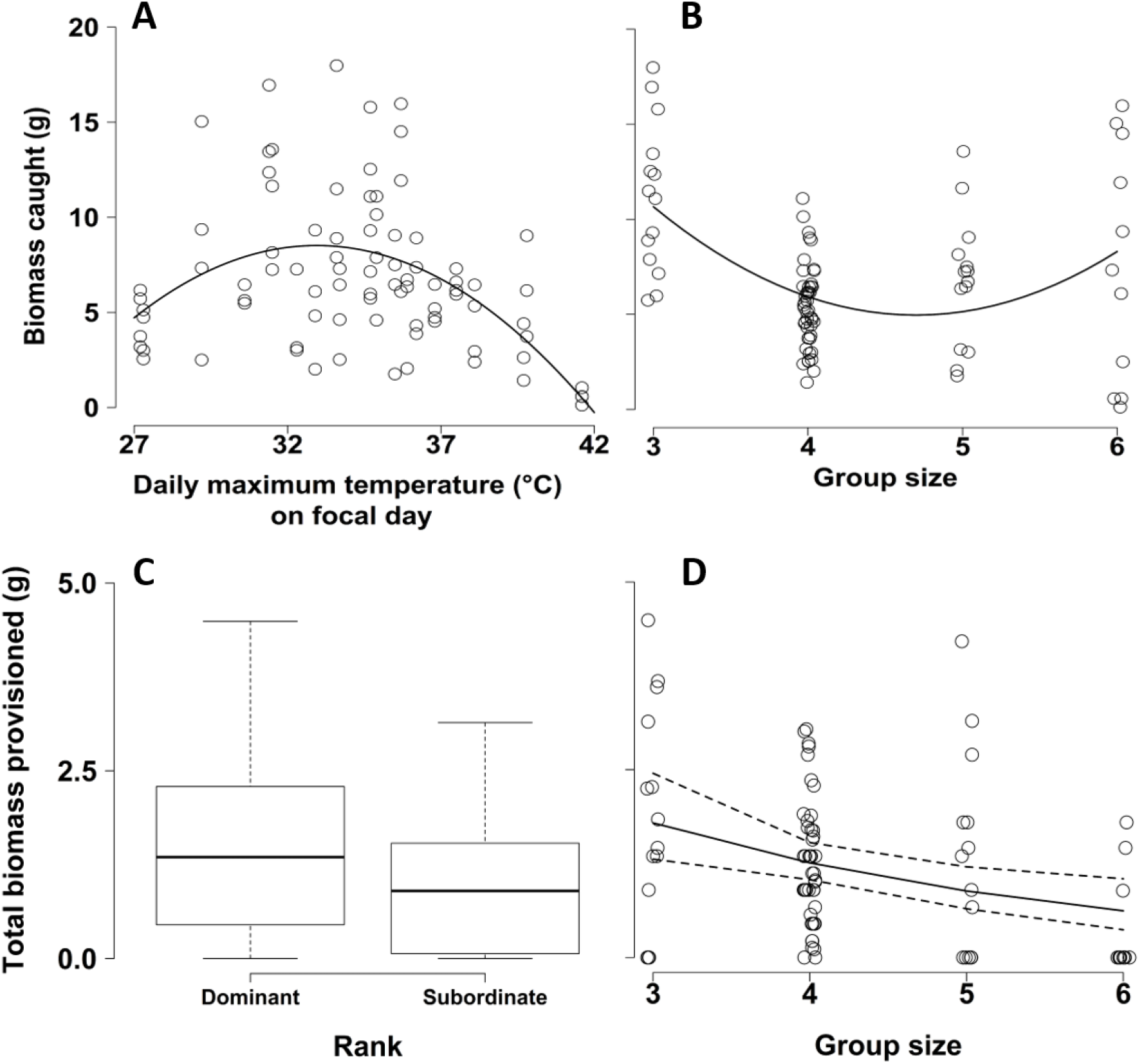
Biomass caught per individual per day as a function of daily maximum temperature (A, in °C) and group size (B) on the observation day. Biomass provisioned per individual per day as a function of rank (C) and group size (D) on the observation day.

### Adult behaviour (nestling age day 7-9)

We observed nest attendance behaviour in 54.3% of focals, and birds spent on average 11 ± 23% of their time attending the nest per focal day. However, the total proportion of time individuals spent attending the nest did not vary significantly with any of the included predictor variables (Table 1). Adults foraged at least once in 92.6% of focals and overall foraged for 53 ± 29% of the time per focal day. Proportion of time spent foraging tended to be higher for males than females and was significantly lower in larger groups and after rain. Proportion of time spent foraging was not influenced by rank, brood size, or T_max_ (Table 1). Birds spent on average 28 ± 25% time resting per focal day. The proportion of time spent resting was significantly higher on hot days, after rain, and in larger groups, but was not influenced by sex, brood size, or rank (Table 1).

**Table 1:** GLMM with Penalised Quasi-Likelihood (glmmPQL) model outputs for factors influencing proportion of time spent foraging, resting, and attending the nest by pied babblers with nestlings aged 7-9 days after hatching. Models fitted to data from 593 focal observations on 62 different individuals from 13 groups collected over 88 days. Significant terms (P < 0.05) are highlighted in bold. Random term: brood identity

| Model | Parameters | estimate | SE | t-value | P-value |
| --- | --- | --- | --- | --- | --- |
| Proportion time spent foraging | *Intercept* | -0.047 | 0.109 | -0.437 | 0.664 |
|  | Brood size | 0.157 | 0.092 | 1.701 | 0.105 |
|  | **Group size** | **-0.275** | **0.090** | **-3.049** | **0.007** |
|  | Maximum temperature | -0.114 | 0.094 | -1.221 | 0.227 |
|  | **Rainfall two months prior** | **-0.322** | **0.093** | **-3.476** | **0.003** |
|  | Rank | -0.013 | 0.102 | -0.132 | 0.895 |
|  | Sex (Male) | 0.208 | 0.105 | 1.986 | 0.051 |
| Proportion time spent resting | *Intercept* | **-0.965** | **0.134** | **-7.226** | **0.000** |
|  | Brood size | -0.176 | 0.104 | -1.689 | 0.108 |
|  | **Group size** | **0.364** | **0.100** | **3.625** | **0.002** |
|  | **Maximum temperature** | **0.247** | **0.109** | **2.259** | **0.027** |
|  | **Rainfall two months prior** | **0.437** | **0.104** | **4.222** | **0.001** |
|  | Rank | 0.154 | 0.133 | 1.153 | 0.254 |
|  | Sex (Male) | -0.049 | 0.138 | -0.356 | 0.723 |
| Proportion time spent attending the nest | *Intercept* | **-1.768** | **0.168** | **-10.507** | **0.000** |
|  | Brood size | -0.019 | 0.115 | -0.164 | 0.872 |
|  | Maximum temperature | -0.208 | 0.117 | -1.781 | 0.079 |
|  | Group size | -0.059 | 0.119 | -0.496 | 0.626 |
|  | Rainfall two months prior | -0.219 | 0.127 | -1.722 | 0.101 |
|  | Rank | -0.284 | 0.223 | -1.275 | 0.207 |
|  | Sex (Male) | -0.403 | 0.226 | -1.779 | 0.080 |

Total biomass caught per observation day averaged 7.0 ± 4.4 g per bird per focal day (*n* = 108 focal days, range: 0.0–20.3 g). After averaging the two top models (combined weight = 0.784), total biomass caught increased with increasing T_max_ until ~32.3°C, after which it declined with increasing T_max_ (*Z* = 3.980, *P* < 0.001; Table 2, Fig. 4A). The effect of group size was also quadratic: birds in intermediate-sized groups caught less biomass per day than birds in larger and smaller groups (*Z* = 3.621, *P* < 0.001; Table 2, Fig. 4B). We found no evidence that biomass caught per day was influenced by sex, rank, brood size, or rainfall (see Table S4 for full model selection outputs).

**Table 2:**
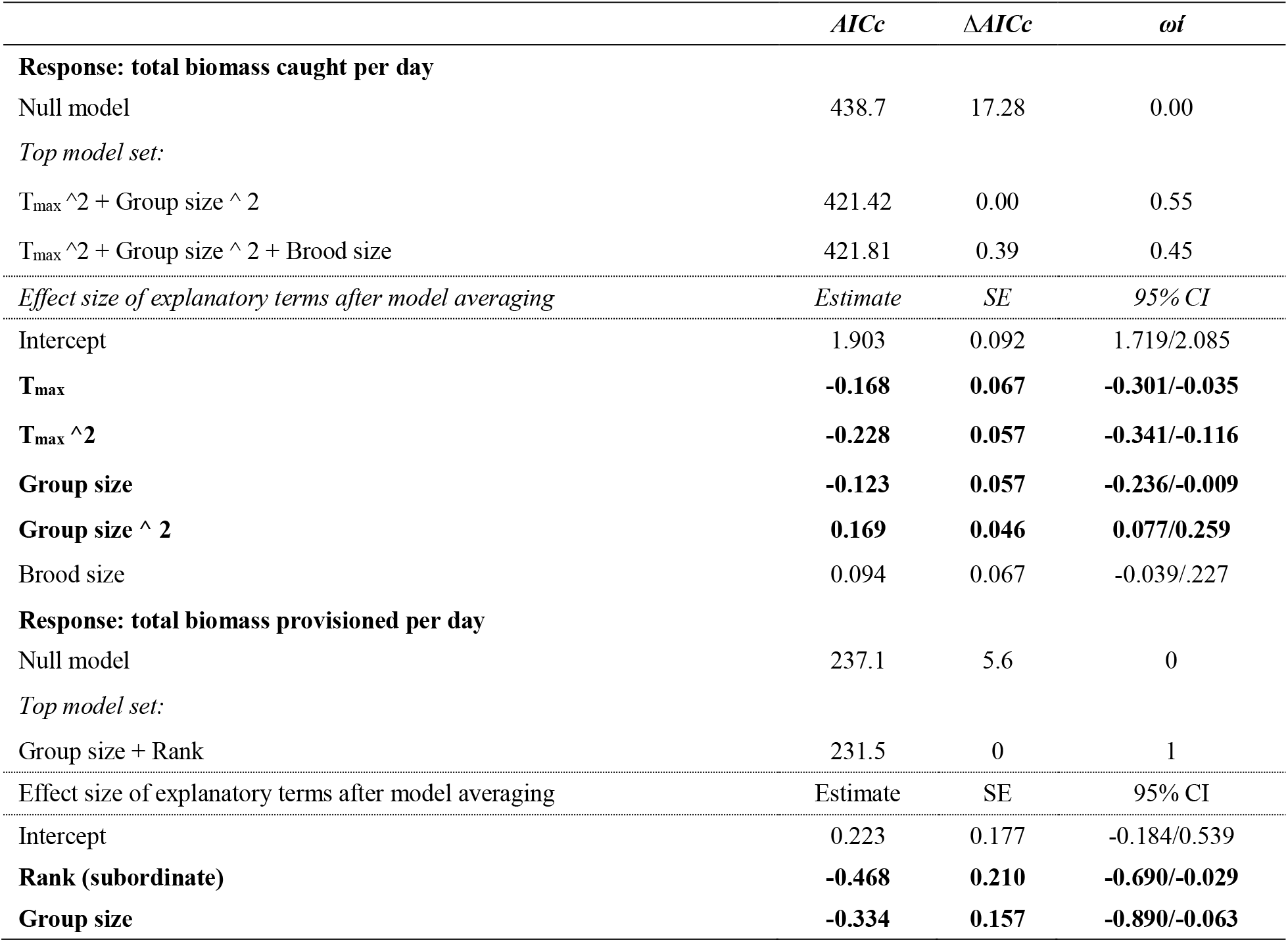
Top model sets for factors influencing total biomass caught per day and total biomass provisioned per day to nestlings 7 to 9 days after hatching. Model averaging was implemented for models with ΔAICc < 2 of the ‘best-fit’ model. Significant terms after model averaging are shown in bold. Data from 56 different individuals from 23 nests by 13 groups over 84 focal days (see Table S4 and S5 for full model selection outputs).

Total biomass provisioned to the nest per observation day averaged 1.2 ± 1.1 g per bird per focal day (*n* = 108 focal days, range: 0.0–4.5 g). Rank and group size were the most parsimonious predictors of variation in biomass provisioned to the nest (the single best-fit model had a model weight of 0.590; Table 2). Subordinate individuals provisioned significantly less (1.0 ± 0.9 g per focal day) than dominant individuals (1.4 ± 1.3 g; *Z* = −2.228, *p* = 0.026; Fig. 4C). Individuals in larger groups provisioned less than those in smaller groups (*Z* = −2.133, *p* = 0.033; Fig. 4D). We found no evidence that biomass provisioned per day was influenced by temperature, brood size, rainfall, or sex (see Table S5 for full model selection outputs).

## Discussion

We investigated the influence of temperature and rainfall on survival of young over the development period from hatching to fledging, in a cooperatively breeding bird endemic to an arid environment (the southern Kalahari Desert) heavily affected by climate change (Kruger and Sekele 2013; van Wilgen et al. 2016; Bourne, Cunningham, et al. 2020a; Bourne, Cunningham, et al. 2020c). We found that heavier nestlings were significantly more likely to fledge, consistent with previous studies showing that smaller size in nestlings correlates with reduced survival (Magrath 1991; Schwagmeyer and Mock 2008), and this effect was detectable in nestlings as young as five days old. Piecewise SEM showed a strong negative influence of high temperatures during the day on nestling body mass in the evening both directly (in both d5 and d11 nestlings) and indirectly via impacts on adult provisioning rates (in d11 nestlings only). Multivariate analysis showed that this result was mirrored by, and potentially explained by, compromised diurnal (12hr) mass gain in both d5 and d11 nestlings on hotter days. Rainfall and group size positively influenced the number of provisioning visits by adults to both d5 and d11 nestlings but did not directly influence nestling structural size in the evening (tarsus and wing length) or daily growth rates. Individual adult pied babblers maintained the overall proportion of time spent attending the nest as well as the total biomass provisioned to nestlings at high temperatures, despite reducing the total number of provisioning visits to the nest overall, and despite declines in the biomass of prey caught per individual as temperatures increased.

That nestling growth was compromised over the course of a single hot day, and that the impacts of reduced body mass on ultimate fledging probability were evident in nestlings as young as five days old, suggests that isolated hot days are likely to be detrimental to nestling survival regardless of whether or not they occur as part of a heat wave. This is consistent with work on other Kalahari species, suggesting that single hot days, even fairly early in the nestling period, may continue to affect nestlings up until fledging age (Cunningham et al. 2013). The effect of temperature on tarsus and wing growth was smaller than for body mass, suggesting that pied babbler nestlings prioritise limb growth over mass gain under challenging conditions. This is expected because individual survival in birds depends strongly on physical traits such as wing length (Naef-Daenzer and Grüebler 2016; Martin et al. 2018). Longer wings allow for improved mobility (Jones et al. 2017), better competitive and predator avoidance abilities (Greño et al. 2008), synchronous fledging (Nilsson and Svensson 1996), and reduced mortality of juveniles (Martin et al. 2018).

Piecewise SEM suggested that direct effects of temperature on evening mass of both d5 and d11 nestlings were more important than indirect effects via adult provisioning rates (also see van de Ven, McKechnie, et al., 2020), despite the fact that parental care behaviours mediate effects of weather conditions on nestling growth in other bird species (Weimerskirch et al. 2000; Tremblay et al. 2005; Cunningham et al. 2013). In this study, the number of provisioning visits was only important for predicting evening mass, and thus fledging probability, of d11 nestlings. From the focal behaviour observations of adults in groups with 7- to 9-day-old nestlings, i.e. just before d11 nestling measurements, we learned that adult pied babblers spent a larger proportion of time resting on hotter than cooler days. However, they did not significantly reduce the proportion of time spent foraging or attending the nest to achieve this, suggesting that they instead reduced their time spent on other behaviours (such as vigilance and defending the territory against neighbouring groups). High temperatures negatively affected total biomass caught and the number of provisioning visits adults made to the nest (consistent with Wiley & Ridley 2016), but not the total amount provisioned per bird per day. Thus, birds flew to the nest less frequently but took larger loads each time. This implies that the only concession provisioning adults make at high temperatures is to shift from a rate-maximising strategy (frequent visits to the nest) to an efficiency-maximising strategy (providing a consistent amount of food to nestlings, in terms of total biomass per day, while limiting the number of provisioning flights by adults as temperatures rise) as temperatures increase.

Less frequent provisioning trips on hotter days are likely to help adult birds avoid raising body temperature by flying (Engel et al. 2006), as previously suggested by Wiley & Ridley (2016). However, shifting to an efficiency-maximising strategy appears insufficient to offset the direct effects of high temperatures on nestling body mass, tarsus and wing length, and daily growth rates. Wet biomass intake may therefore need to increase at high temperatures to maintain nestling mass and sustain growth, due to increased nestling demand for water to aid thermoregulation under hot conditions (Salaberria et al. 2014; van de Ven, McKechnie, et al. 2020; Czenze et al. 2020). If chicks become dehydrated, growth could be hampered by poor physiological performance due to costs associated with dehydration and subsequent high body temperatures (Angilletta et al. 2010). If such elevated water demand does exist when hot, our data suggest that provisioning adults might be unable to increase biomass provisioned in order meet this demand. This is because biomass caught declined with increasing temperature above 32.3ºC, indicating poorer foraging success (du Plessis et al. 2012; Cunningham et al. 2015; van de Ven et al. 2019) and suggesting that there was probably less biomass available at high temperatures with which to provision to nestlings (Conrey et al. 2016; Dodson et al. 2016; Mella et al. 2018). Adults may also be constrained in their ability to provision more water-rich food to nestlings at high temperatures due to the increased costs of flight at high air temperatures (Klaassen 1995; Powers et al. 2017) and the need to attend to their own water demands (Bourne, Ridley, et al. 2020; Czenze et al. 2020).

Nestling body mass, tarsus and wing length, and daily growth rates were not affected by group size (also see Wiley & Ridley 2016), suggesting that the benefits of cooperation accrue to adult group members rather than young in this species in terms of nestling growth (Mumme et al. 2015; Savage et al. 2015; Langmore et al. 2016). In keeping with this, adult behaviour was affected by group size. Nestlings received the same level of care across group sizes, but adults in larger groups invested less per individual in raising young than adults in smaller groups: they spent less time foraging and more time resting than individuals in smaller groups, and provisioned less biomass to young per individual on average than those in smaller groups. This adds further support to previous evidence that ‘load-lightening’ occurs in pied babblers (Raihani and Ridley 2008; Ridley and Raihani 2008; Wiley and Ridley 2016).

## Conclusion

High temperatures during the nestling period affected the mass, tarsus length, and survival to fledging of pied babbler nestlings, consistent with prior research on passerines (Greño et al. 2008; Salaberria et al. 2014), including this species (Wiley and Ridley 2016; Bourne, Cunningham, et al. 2020a). Piecewise SEM analysis suggests that the majority of this effect is driven by direct effects of temperature on nestlings, and to a lesser extent by temperature-mediated changes in provisioning rate to older nestlings. Although parental care strategies are flexible in response to both climate and social conditions, these strategies have limits (Cunningham et al. 2013; van de Ven, McKechnie, et al. 2020). Our results suggest that mitigatory actions by provisioning adults (e.g. the behavioural shifts from rate-maximising to efficiency-maximising provisioning strategies documented here) failed to compensate fully for direct effects of high temperatures on nestling growth and development. Repeated exposure to high temperatures during breeding attempts could therefore undermine population replacement via low recruitment of young into the adult breeding population, leading to an increasingly detrimental impact of high temperatures on population persistence over time (Bourne, Cunningham, et al. 2020a; Bourne, Cunningham, et al. 2020b; Bourne, Cunningham, et al. 2020c). This suggests a mechanism by which predicted temperature increases in the Kalahari (MacKellar et al. 2014) could negatively affect population growth and persistence (Cunningham *et al.* 2013, Conradie *et al.* 2019).

We highlight the need to quantify multiple simultaneous factors with direct as well as indirect pathways that influence fledging probabilities, including adult behaviour, investment in parental care, and offspring growth and development, to identify mechanisms by which birds are at risk under global change. Sublethal costs of high temperatures such as those documented here should not be overlooked as they are likely to prove powerful drivers of population decline under climate change (Conradie et al. 2019; Román-Palacios and Wiens 2020; Trisos et al. 2020). Impacts on breeding success appear to operate similarly across multiple species (Cunningham et al. 2013; Salaberria et al. 2014; Cooper et al. 2019; Sharpe et al. 2019; Exposito-Granados et al. 2019 Sep 24; Bourne, Cunningham, et al. 2020a; van de Ven, McKechnie, et al. 2020) and could cause collapses of animal communities (Iknayan and Beissinger 2018; Riddell et al. 2019; Ripple et al. 2019; Rosenberg et al. 2019) long before temperatures reach and begin to regularly exceed lethal limits.

