## Supplementary materials for "Direct and indirect effects of high temperatures on fledging success in a cooperatively breeding bird"

**Title: High temperatures are associated with poor nestling growth and altered parental care and likely limit fledging rates in a cooperatively breeding bird**

Amanda R. Bourne*[1], Amanda R. Ridley[1,2], Claire N. Spottiswoode[1,3], Susan J. Cunningham[1]

[1] FitzPatrick Institute of African Ornithology, DST-NRF Centre of Excellence, University of Cape Town, Private Bag X3, Rondebosch 7701, South Africa

[2] Centre for Evolutionary Biology, School of Biological Sciences, University of Western Australia, Crawley 6009, Australia

[3] Department of Zoology, University of Cambridge, Downing Street, Cambridge CB2 3EJ, UK

#### Supporting Information

While there were time of day effects on foraging behaviour (e.g. foraging success, one-way ANOVA $F_{5,489} = 5.390$, $P < 0.001$), sampling evenly across the day per individual ensured no time of day biases on data collected ($F_{5,489} = 1.283$, $P = 0.269$), such that differences between days could be attributed to factors occurring on that day, rather than being artefacts of the time of day at which data were collected. Data analysed at the scale of the focal day are therefore comparable between birds and between days because we collected the same quantity of data per bird and per session across days.

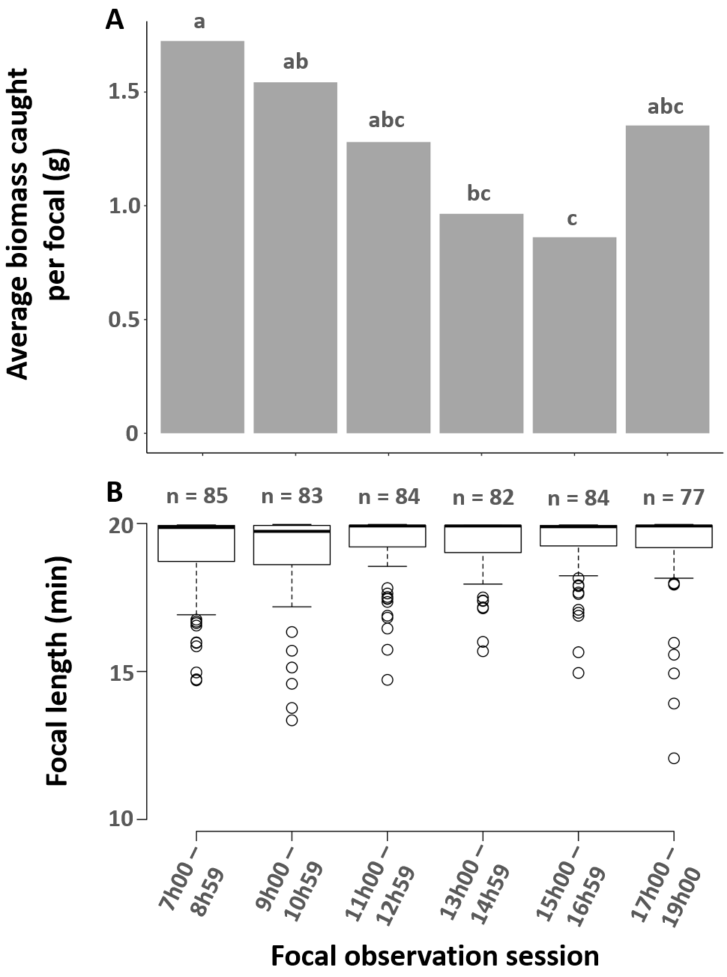


*Figure S1: (A) average biomass caught per focal, and (B) focal length for each focal observation session. Post-hoc comparisons (Tukey HSD) show which focal sessions differed significantly from one another with regards to average biomass caught (A). Because focal data were collected evenly across all times of day, these differences in prey capture rates between focals will not affect comparisons between different days at the scale of whole days.*

#### Path analysis diagrams for nestling size

#### Tarsus length, d5

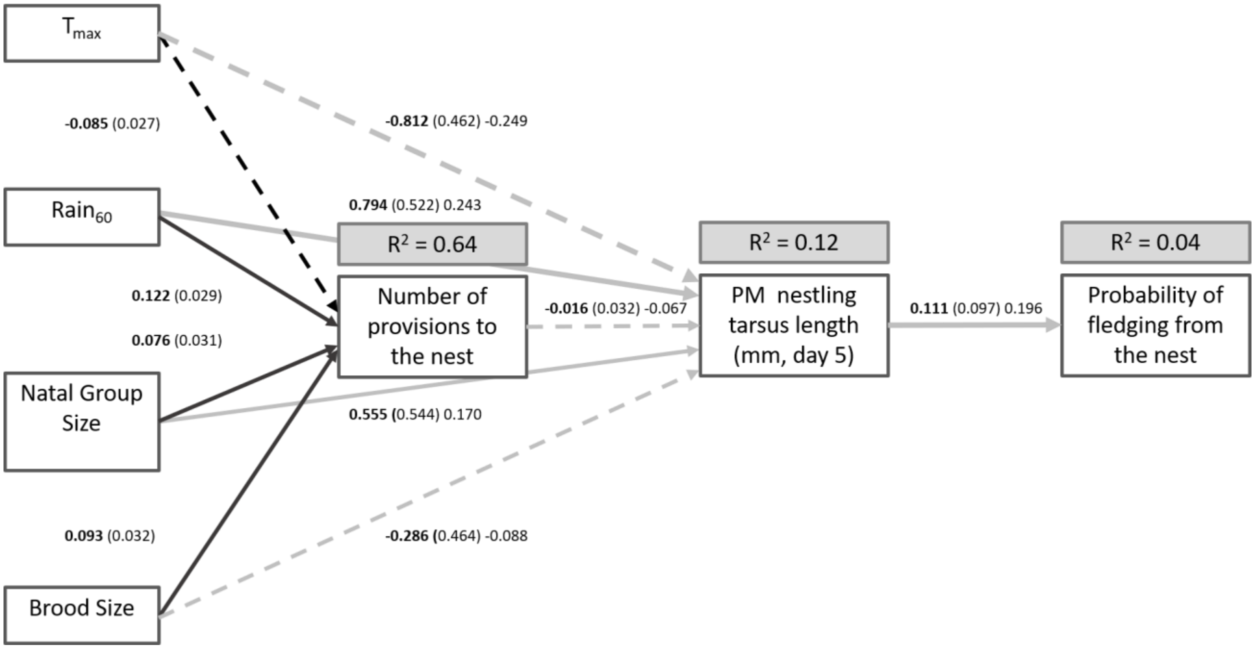


*Figure S2: Piecewise SEM exploring the effects of environmental factors (temperature and rainfall), group size, and brood size on individual probabilities of fledging via number of provisions and nestling tarsus length (evening) 5 days after hatching. Boxes represent measured variables. Arrows represent unidirectional relationships among variables. Solid arrows denote positive relationships, dashed arrows negative relationships. Non-significant paths are grey. Path coefficients are shown in bold, followed by standard errors in parentheses. Standardised coefficients are shown after standard errors for all pathways except those in the Poisson-distributed model with number of provisions to the nest as the response variable, for which standardised estimates could not be calculated. Path thickness has been scaled relative to the absolute magnitude of the standardised estimates, where available, such that stronger effects have thicker arrows.* $R^2$ *for component models are given in the grey boxes above response variables. Model fit: Fisher's C = 14.83, df = 10, p = 0.139.*

#### Wing length, d5

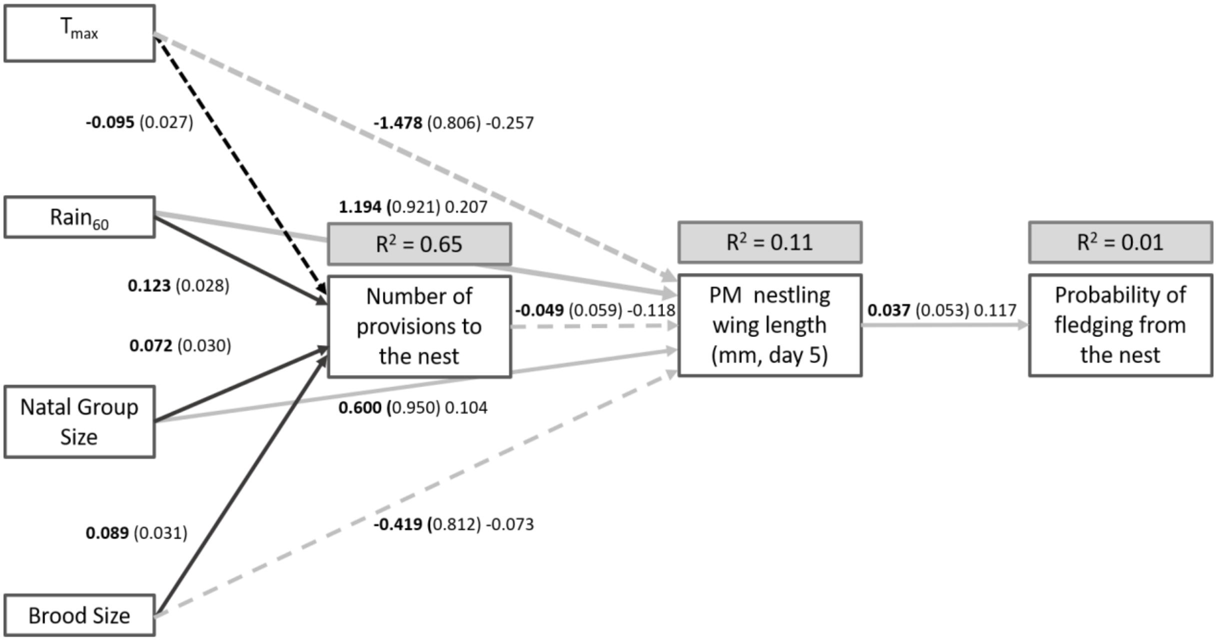


*Figure S3: Piecewise SEM exploring the effects of environmental factors (temperature and rainfall), group size, and brood size on individual probabilities of fledging via number of provisions and nestling wing length (evening) 5 days after hatching. Boxes represent measured variables. Arrows represent unidirectional relationships among variables. Solid arrows denote positive relationships, dashed arrows negative relationships. Non-significant paths are grey. Path coefficients are shown in bold, followed by standard errors in parentheses. Standardised coefficients are shown after standard errors for all pathways except those in the Poisson-distributed model with number of provisions to the nest as the response variable, for which standardised estimates could not be calculated. Path thickness has been scaled relative to the absolute magnitude of the standardised estimates, where available, such that stronger effects have thicker arrows. R2 for component models are given in the grey boxes above response variables. Model fit: Fisher's C = 15.08, df = 10, p = 0.129.*

#### Tarsus length, d11

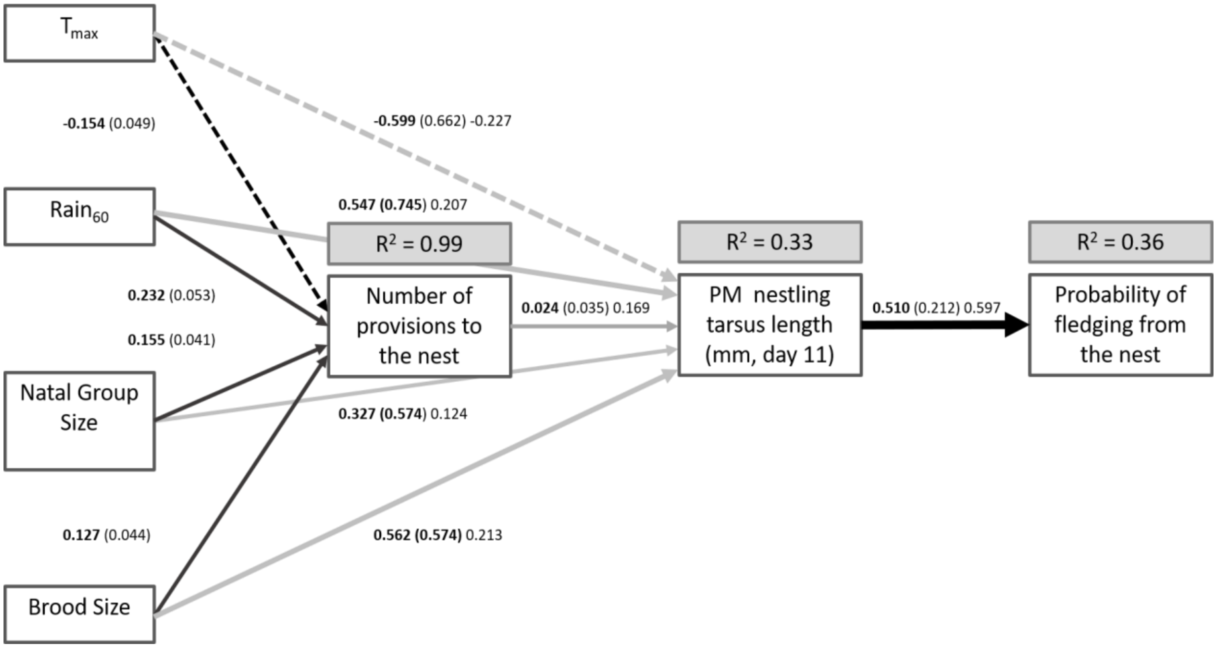


*Figure S4: Piecewise SEM exploring the effects of environmental factors (temperature and rainfall), group size, and brood size on individual probabilities of fledging via number of provisions and nestling tarsus length (evening) 11 days after hatching. Boxes represent measured variables. Arrows represent unidirectional relationships among variables. Solid arrows denote positive relationships, dashed arrows negative relationships. Non-significant paths are grey. Path coefficients are shown in bold, followed by standard errors in parentheses. Standardised coefficients are shown after standard errors for all pathways except those in the Poisson-distributed model with number of provisions to the nest as the response variable, for which standardised estimates could not be calculated. Path thickness has been scaled relative to the absolute magnitude of the standardised estimates, where available, such that stronger effects have thicker arrows.* $R^2$ *for component models are given in the grey boxes above response variables. Model fit: Fisher's C = 10.13, df = 10, p = 0.429.*

#### Wing length, d11

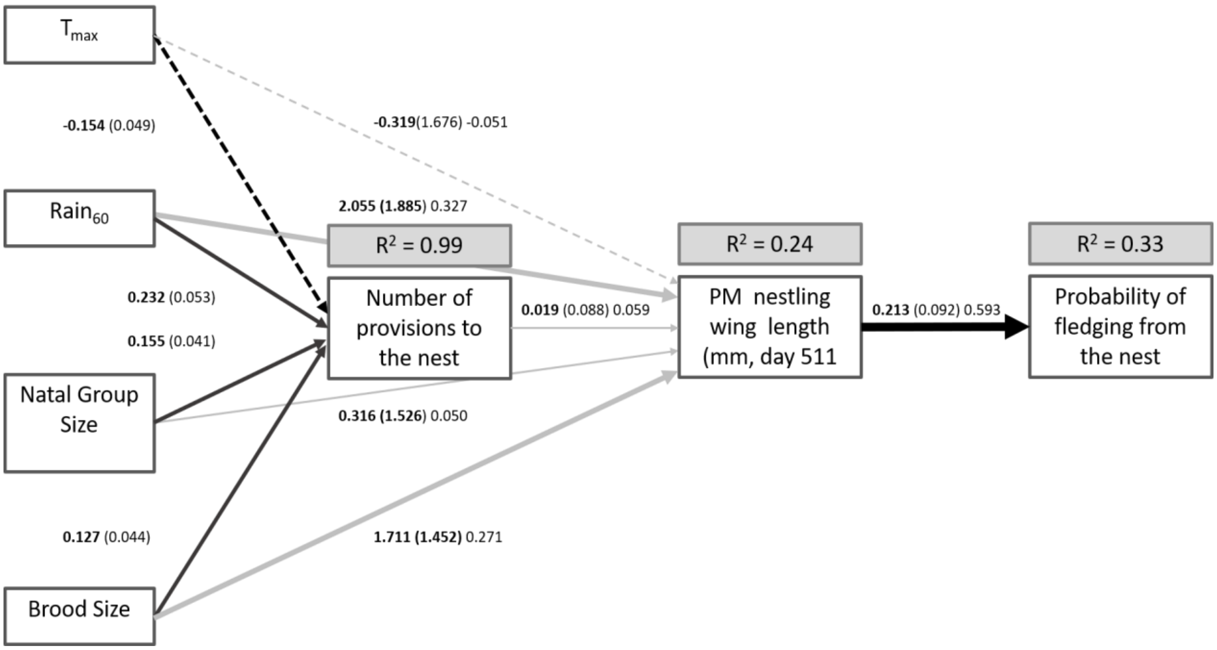


*Figure S5: Piecewise SEM exploring the effects of environmental factors (temperature and rainfall), group size, and brood size on individual probabilities of fledging via number of provisions and nestling wing length (evening) 11 days after hatching. Boxes represent measured variables. Arrows represent unidirectional relationships among variables. Solid arrows denote positive relationships, dashed arrows negative relationships. Non-significant paths are grey. Path coefficients are shown in bold, followed by standard errors in parentheses. Standardised coefficients are shown after standard errors for all pathways except those in the Poisson-distributed model with number of provisions to the nest as the response variable, for which standardised estimates could not be calculated. Path thickness has been scaled relative to the absolute magnitude of the standardised estimates, where available, such that stronger effects have thicker arrows.* $R^2$ *for component models are given in the grey boxes above response variables. Model fit: Fisher's C = 13.45, df = 10, p = 0.200.*

### Model output tables

*Table S1: Full LMM model outputs for analyses of nestling daily mass change at 5 days old. Data from 93 nestlings weighed am and pm at 5 days old, from 37 breeding attempts by 19 different groups over 3 breeding seasons. Random terms: Brood identity.*

| ***Model Term*** | ***AICc*** | ***ΔAICc*** | ***weight*** |
|---|---|---|---|
| *Null model* | *726.9* | *5.03* | *0.052* |
| Brood size | 725.6 | 3.7 | 0.102 |
| $T_{max}$ | 721.9 | 0 | 0.645 |
| $Rain_{60}$ | 725.4 | 3.56 | 0.109 |
| Group size | 725.8 | 3.88 | 0.092 |
| ***Top models*** | ***AICc*** | ***ΔAICc*** | ***weight*** |
| Null model | 726.9 | 5 | 0 |
| $T_{max}$ | 721.9 | 0 | 1 |
| Effect size of explanatory terms after model averaging | Estimate | SE | 95% CI |
| Intercept | 23.737 | 1.944 | 19.927/27.534 |
| **$T_{max}$** | **-4.043** | **1.986** | **-7.922/-0.151** |

*Table S2: Full LMM model outputs for analyses of nestling daily mass change at 11 days old. Data from 77 nestlings weighed am and pm at 11 days old, from 34 breeding attempts by 18 different groups over 3 breeding seasons. Random terms: Brood identity.*

| ***Model Term*** | ***AICc*** | ***ΔAICc*** | ***weight*** |
|---|---|---|---|
| Null model | 540.2 | 29.84 | 0 |
| Brood size | 539.4 | 29.08 | 0 |
| $T_{max}$ | 510.4 | 0 | 1 |
| $Rain_{60}$ | 539.8 | 29.42 | 0 |
| Group size | 539.3 | 28.95 | 0 |
| ***Top models*** | ***AICc*** | ***ΔAICc*** | ***weight*** |
| Null model | 540.2 | 29.8 | 0 |
| $T_{max}$ | 510.4 | 0 | 1 |
| Effect size of explanatory terms after model averaging | Estimate | SE | 95% CI |
| Intercept | 6.697 | 1.077 | 4.590/8/806 |
| **$T_{max}$** | **-7.028** | **1.122** | **-9.316/-4.834** |

*Table S3: Full LMM model outputs for analyses of nestling daily tarsus length change at 11 days old. Data from 77 nestlings weighed am and pm at 11 days old, from 34 breeding attempts by 18 different groups over 3 breeding seasons. Random terms: Brood identity.*

| ***Model Term*** | ***AICc*** | ***ΔAICc*** | ***weight*** |
|---|---|---|---|
| Null model | 346.6 | 4.52 | 0.081 |
| Brood size | 347.3 | 5.26 | 0.056 |
| $T_{max}$ | 342.1 | 0 | 0.772 |
| $Rain_{60}$ | 349.3 | 7.25 | 0.021 |
| Group size | 346.8 | 4.77 | 0.071 |
| ***Top models*** | ***AICc*** | ***ΔAICc*** | ***weight*** |
| Null model | 346.6 | 4.5 | 0 |
| $T_{max}$ | 342.1 | 0 | 1 |
| Effect size of explanatory terms after model averaging | Estimate | SE | 95% CI |
| Intercept | 2.945 | 0.279 | 2.401/3.501 |
| **$T_{max}$** | **-0.804** | **0.287** | **-1.381/-0.245** |

*Table S4: Full GLMM model outputs for analyses of total biomass caught. Data from 84 focal days collected on 56 different individuals at 22 breeding attempts by 13 different groups over 3 breeding seasons. Random terms: Brood identity.*

| ***Model Term*** | ***AICc*** | ***ΔAICc*** | ***weight*** |
|---|---|---|---|
| Null model | 438.7 | 17.33 | 0 |
| Sex | 438.6 | 17.16 | 0 |
| Rank | 440.9 | 19.45 | 0 |
| Brood size | 434.6 | 13.21 | 0.001 |
| $T_{max}$ | 438.6 | 17.17 | 0 |
| $T_{max}$ ^ 2 | 426.3 | 4.84 | 0.038 |
| $Rain_{60}$ | 435.6 | 14.17 | 0 |
| Group size | 438.8 | 17.4 | 0 |
| Group size ^ 2 | 435.7 | 14.26 | 0 |
| Brood size + $Rain_{60}$ | 431.7 | 10.25 | 0.003 |
| $T_{max}$ ^ 2 + Brood size | 427 | 5.58 | 0.026 |
| Group size ^ 2 + $Rain_{60}$ | 435.3 | 13.92 | 0 |
| $T_{max}$ ^ 2 + $Rain_{60}$ | 427.9 | 6.44 | 0.017 |
| $T_{max}$ ^2 + Group size ^ 2 | 421.4 | 0 | 0.43 |
| Group size ^ 2 + Brood size | 430 | 8.61 | 0.006 |
| $T_{max}$ ^ 2 + $Rain_{60}$ + Brood size | 428.3 | 6.85 | 0.014 |
| $T_{max}$ ^ 2 + Brood size + Group size ^2 | 421.8 | 0.39 | 0.354 |
| $T_{max}$ ^ 2 + $Rain_{60}$ + Brood size + Group size ^2 | 424.2 | 2.74 | 0.11 |

*Table S5: Full GLMM model outputs for analyses of total biomass provisioned. Data from 84 focal days collected on 56 different individuals at 22 breeding attempts by 13 different groups over 3 breeding seasons. Random terms: Brood identity.*

| ***Model Term*** | ***AICc*** | ***ΔAICc*** | ***weight*** |
|---|---|---|---|
| Null model | 237.1 | 5.61 | 0.036 |
| Sex | 239.1 | 7.66 | 0.013 |
| Rank | 233.8 | 2.36 | 0.181 |
| Brood size | 238.5 | 7.01 | 0.018 |
| $T_{max}$ | 239.2 | 7.72 | 0.012 |
| $Rain_{60}$ | 238.7 | 7.24 | 0.016 |
| Group size | 234.4 | 2.95 | 0.135 |
| Group size + Rank | 231.5 | 0 | 0.59 |
| ***Top models*** | ***AICc*** | ***ΔAICc*** | ***weight*** |
| Basic (1 + (1\|NestCode) | 237.1 | 5.6 | 0 |
| Group size + Rank | 231.5 | 0 | 1 |
| Effect size of explanatory terms after model averaging | Estimate | SE | 95% CI |
| Intercept | 0.223 | 0.177 | -0.184/0.539 |
| **Rank** | **-0.468** | **0.210** | **-0.690/-0.029** |
| **Group size** | **-0.334** | **0.157** | **-0.890/-0.063** |
